# Clinically Relevant Pathogens on Surfaces Display Differences in Survival and Transcriptomic Response in Relation to Probiotic and Traditional Cleaning Strategies

**DOI:** 10.1101/2022.01.11.475867

**Authors:** Jinglin Hu, Weitao Shuai, Jack T. Sumner, Anahid A. Moghadam, Erica M. Hartmann

## Abstract

Indoor surfaces are paradoxically presumed to be both colonized by pathogens, necessitating disinfection, and “microbial wastelands.” In these resource-poor, dry environments, competition and decay are thought to be important drivers of microbial community composition. However, the relative contributions of these two processes have not been specifically evaluated. To bridge this knowledge gap, we used microcosms to specifically evaluate whether interspecies interactions occur on surfaces. We combined transcriptomics and traditional microbiology techniques to investigate whether competition occurred between two clinically important pathogens, *Acinetobacter baumannii* and *Klebsiella pneumoniae*, and a probiotic cleaner containing a consortium of *Bacillus* species. Probiotic cleaning seeks to take advantage of ecological principles such as competitive exclusion, thus using benign microorganisms to inhibit viable pathogens, but there is limited evidence that competitive exclusion in fact occurs in environments of interest (i.e., indoor surfaces). Our results indicate that competition in this setting has a negligible impact on community composition but may influence the functions expressed by active organisms. Although *Bacillus* spp. remained viable on surfaces for an extended period of time after application, viable colony forming units (CFUs) of *A. baumannii* recovered following exposure to a chemical-based detergent with and without *Bacillus* spp. showed no statistical difference. Similarly, for *K. pneumoniae*, there were small statistical differences in CFUs between cleaning scenarios with or without *Bacillus* spp. in the chemical-based detergent. The transcriptome of *A. baumannii* with and without *Bacillus* spp. exposure shared a high degree of similarity in overall gene expression, but the transcriptome of *K. pneumoniae* differed in overall gene expression, including reduced response in genes related to antimicrobial resistance. Together, these results highlight the need to fully understand the underlying biological and ecological mechanisms for community assembly and function on indoor surfaces, as well as having practical implications for cleaning and disinfection strategies for infection prevention.

## 1. Introduction

Indoor surfaces harbor pathogens and mediate the transmission of infectious disease. At the same time, these environments have been called a “microbial wasteland” where microbes are “dead, dying, or dormant.” Following this model, surface-associated microbial communities are almost entirely driven by the opposing forces of dispersal and death. Surface cleaning and disinfection practices seek to promote microbial cell death, thereby preventing infection, especially in healthcare settings. While cleaning using antimicrobials can effectively reduce microbial load on contaminated surfaces in health care facilities^1^, evidence of their effects on preventing disease transmission and reducing healthcare-acquired infections remain limited^2^. Moreover, because antimicrobials can cause both acute and chronic health impacts^3^ and lead to antimicrobial resistance^4–6^, there is a strong drive to develop alternative cleaning strategies.

Probiotic cleaning seeks to take advantage of competitive exclusion and other ecological principles to inhibit viable pathogens on surfaces^4, 7^. Members of *Bacillus*, *Bifidobacterium*, *Lactobacillus*, *Rhodopseudomonas*, and *Saccharomyces* are the most frequently used microorganisms in these microbial based cleaning products^8^. While probiotic organisms may persist on surfaces, competitive exclusion is not compatible with the “wasteland” model, which stipulates no growth or metabolic activity. Previous studies of probiotic cleaners have shown that probiotic cleaning decreases the recovery of cultivable bacterial pathogens^9, 7^, and even inactivates viruses^10^. Competitive exclusion is an unlikely mechanism of action against viruses, but even for bacteria there is limited direct evidence documenting competitive exclusion^11^. Rather, culture-based results support the hypothesis that probiotic *Bacillus* outcompete pathogens in some environmental conditions but are in fact out-competed by resident organisms when the latter are present in biofilms^11^. Moreover, there is a dearth of molecular evidence supporting whether surviving probiotic organisms and healthcare-associated pathogens are engaged in activities such as nutrient scavenging, stress response, or microbe-microbe interactions^11^. To this end, studying gene expression levels instead of cultivable colonies or gene presence in both pathogens and probiotic microorganisms is necessary to unravel the mechanisms of probiotic cleaning. Transcriptomics allows quantitative determination of changes in regulation of genes and pathways of the microorganisms upon application of probiotic cleaners under different conditions, revealing changes in function and metabolic activity. Despite these advantages, the application of transcriptomics in indoor surface microbiome studies has been sparse^12^. Current indoor surface microbiome studies only focus on taxonomic profiling and functional potentials that recovered from (meta)genomics analysis where the importance of gene expression levels is overlooked or yet to be addressed^13, 14^. Transcriptomics analysis from this study can provide insights on the response of pathogens to tested cleaning scenarios on indoor surfaces.

*Acinetobacter baumannii* and *Klebsiella pneumoniae* have been recognized as a major source of healthcare-associated infections associated with high morbidity and mortality rates due to their resistance to many antibiotics including carbapenems^15, 16^. These organisms not only persist on surfaces for up to several months^17^ but are commonly found in healthcare environments, especially in low- and middle-income countries. *K. pneumoniae* have been found on surfaces in hospitals in Brazil^16, 15^, Tunisia^17^, Russia^18^, China^19^ and Ghana^20^ *A. baumannii* was found on surfaces in hospitals in Brazil^21^, Turkey^21^, Saudi Arabia^22^ and on the surfaces of medical devices in Iran^23^. It was also found on surfaces in buildings surrounding hospitals in the US^24^. Both *A. baumannii* and *K. pneumoniae* were found on surfaces in neonatal intensive care units in Egypt^25^ and hospitals in Algeria^26^. While these organisms likely reach surfaces via dispersion from human occupants, they can also be transferred back to humans, particularly the very vulnerable. *K. pneumoniae* persists in neonatal intensive care units and transfers to neonates’ gut^27, 28^ and possibly also the nares^29^. A high degree of relatedness has been observed between genotypes of *A. baumannii* from patients and their surrounding rooms^30^, and bidirectional transfer of *A. baumannii* between the environment and adult patients has also been observed^31^. Notably in this last study, *A. baumannii* then persisted in the environment despite terminal disinfection^31^. More concerning than individual transfer events or colonization, surface-associated organisms can cause or exacerbate outbreaks within a hospital. For example, surface-associated *K. pneumoniae* was associated with an outbreak in Vietnam^32^, and *A. baumannii* on bed railings was associated with an outbreak in Saudi Arabia^33^.

To understand how these organisms survive on surfaces, sometimes in defiance of cleaning efforts, we sought to observe changes in gene expression in *A. baumannii* and *K. pneumoniae* on surfaces with and without cleaning. Sequencing of low biomass samples collected from the built environment often suffers from a low signal-to-noise ratio^34^. To obtain a fundamental mechanistic understanding of indoor cleaning strategies and their impacts on microbial survival and interactions, we utilized microcosm chambers containing stainless steel sheets to simulate a simplified built environment with greater than usual microbial load. Because antimicrobial resistance and hospital-acquired infections disproportionately affect low- and middle-income countries, many of which are located in tropical climates, we also evaluated the effects of elevated temperature and relative humidity (RH). To evaluate general ecological phenomena, and particularly competitive exclusion, governing survival in mixed communities, we further exposed these organisms to a probiotic cleaner containing several strains of *Bacillus*.

## 2. Materials and Method

### 2.1. Microcosm set-up

A commercially available All-Purpose Probiotic Cleaner (referred to as probiotic cleaner herein) was obtained from Graz, Austria. Clinical isolates of *A. baumannii* (ABBL18) and *K. pneumoniae* (CRE231) were provided by Dr. Alan Hauser, Northwestern University Feinberg School of Medicine. To simulate a typical high-touch surface in a healthcare environment, 24G stainless steel sheets were cut into 2”x2” squares. *A. baumannii* and *K. pneumonia*e isolates were grown in 40 mL tryptic soy broth (TSB) for 23 hours at 30°C with continuous shaking to its mid log phase. The probiotic cleaner was filtered through a 0.22 µm syringe filter to create an Cleaner Only control without probiotic materials. Sterility of the Cleaner Only control was assessed by plating on tryptic soy agar (TSA). *Bacillus* spp. were germinated by mixing 0.1% probiotic cleaner with TSB for use as a model Germinated *Bacillus* control where cells were vegetative. This culture was incubated at 25°C for 23 hours with continuous shaking. Cells (*Bacillus* spp., *A. baumannii* and *K. pneumoniae*) were harvested and washed in phosphate-buffered saline (PBS) before inoculation. A total of six testing scenarios for each pathogen (twelve groups in total) were specified as shown in Table 1:

**Table 1.** Testing scenarios and their experimental group names for each pathogen (i.e., *A. baumannii* and *K. pneumoniae)* consisting of four cleaning conditions, and two temperature and humidity conditions. Cleaning conditions include probiotic-containing and probiotic-free All-Purpose Probiotic Cleaner solutions, vegetative *Bacillus* spp. as a model probiotic, and a no cleaning (i.e., pathogen alone) control. Ambient and elevated temperature and relative humidity (referred to as Ambient and Humid respectively) conditions were assessed as environmental variables. N/A means the scenario was not tested.

| Cleaning Scenarios | Temperature and humidity conditions |  |
| --- | --- | --- |
|  | 25°C and 20% RH | 37°C and 90% RH |
| Pathogen only | No Cleaner (Ambient) | No Cleaner (Humid) |
| Pathogen + filtered probiotic cleaner | Cleaner Only | N/A |
| Pathogen + probiotic cleaner | Probiotic Cleaner (Ambient) | Probiotic Cleaner (Humid) |
| Pathogen + germinated *Bacillus* | Germinated *Bacillus* | N/A |

Each 2”x2” stainless steel surface was first inoculated with 50 µL *A. baumannii* (corresponding to 8.5×10^7^ CFU) or 50 µL *K. pneumoniae* (corresponding to 1.0×10^8^ CFU). 50 µL sterilized PBS buffer, filtered probiotic cleaner, probiotic cleaner, or vegetative *Bacillus* spp. inoculum was added to create No Cleaner control samples (*A. baumannii* or *K. pneumoniae* only), Cleaner Only groups (chemical cleaning samples), Probiotic Cleaner groups (probiotic cleaning samples), or Germinated *Bacillus* groups (vegetative cleaning samples), respectively. Following the introduction of bacterial inoculum, all surfaces were transferred into autoclaved mason jars (referred to as microcosm chambers) under two treatment conditions: 1) ambient temperature (25°C) and relative humidity (20%), and 2) elevated temperature (37°C) and relative humidity (>90%) with 10 mL of autoclaved milliQ water as in^13^. The relative humidity was measured using Fisherbrand™ Traceable™ Humidity-On-A-Card™ Humidity Monitor (Waltham, MA, USA). Each microcosm chamber contained one surface sample. All microcosm chambers were stored in incubators maintained at 25°C or 37°C. Samples were sacrificed and biomass recovered by swabbing at three specific timepoints: 3 hours, 24 hours, and 72 hours following the inoculation and cleaning. Triplicate samples were prepared for each combination of cleaning, timepoint, and incubation conditions (Figure 1).

**Figure 1.**
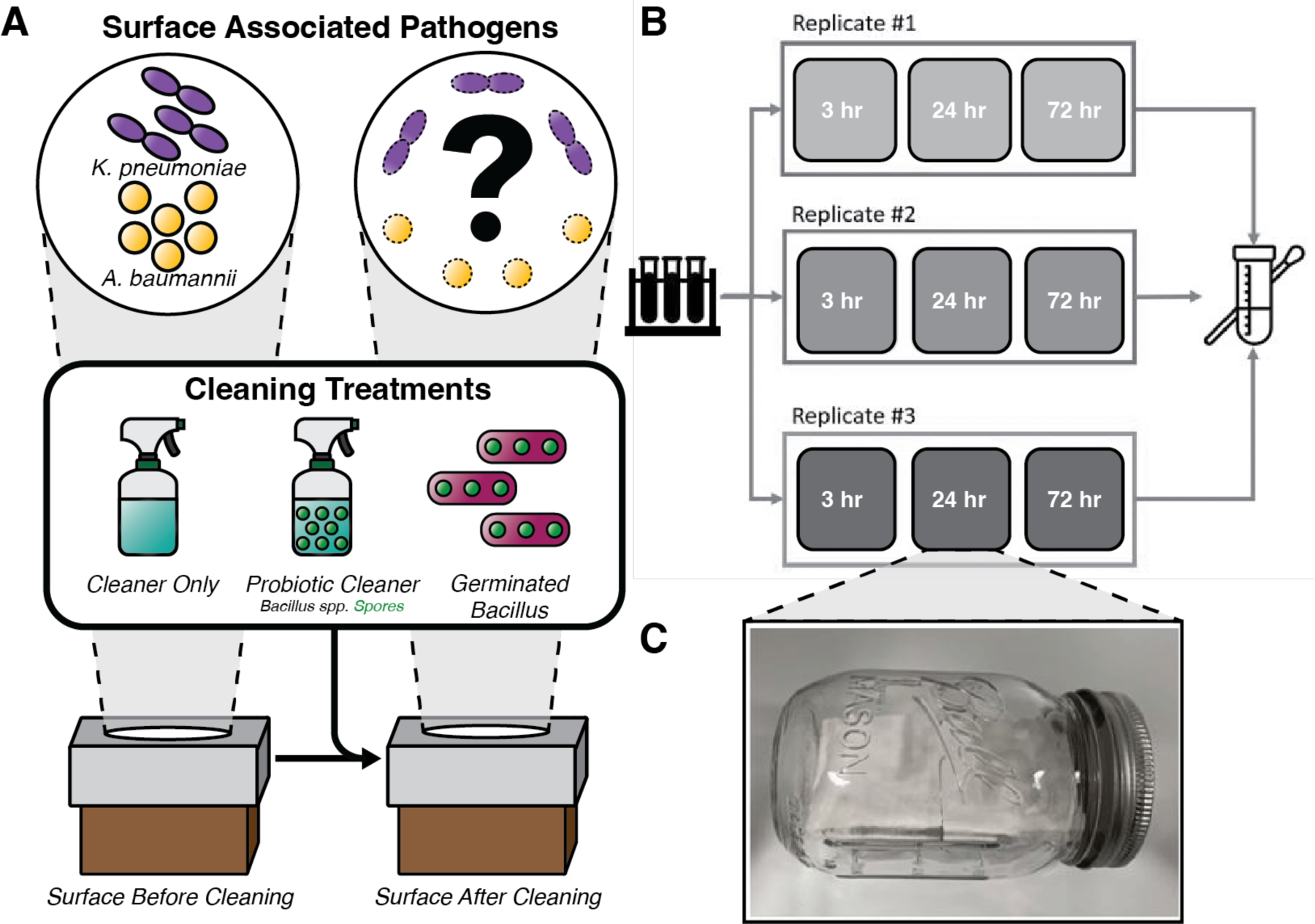
General overview of the biological question (A). Microcosm experimental design (B) with four types of inocula containing pathogen (*A. baumannii* or *K. pneumoniae*) and *Bacillus* spp. and sealed sterile microcosm chamber maintained under ambient temperature and relative humidity (C). The elevated temperature and humidity condition was maintained by adding 10 mL of sterilized miliQ water into the sealed chamber.

### 2.2. Swab sample collection

Swab samples were collected through a combination of dry and wet swabbing as performed previously^13^. Each surface was first dry swabbed from left to right for seven times and the swab was swirled into an aliquoted PBS buffer or RNAprotect Bacteria Reagent (Qiagen), for colony forming units (CFUs) enumeration and RNA extraction, respectively. Surfaces were then wet swabbed for a total of 20 s, including 5 seconds of rinse and rewet. Finally, the swabs were left in 15 mL conical tubes and vortexed for 10 seconds. For total enumeration, 100 µL swab suspension along with its dilutions were spread onto TSA. Additionally, the same volume of swab suspension and dilutions were spread onto CHROM *A. baumannii* selective agar or on MacConkey agar (as *K. pneumoniae* selective media^35^). Under the circumstances where *Bacillus* were introduced through Probiotic Cleaner and Germinated *Bacillus* cleaning, total CFUs were quantified using TSA. All plates were incubated at 37°C. Numbers of CFUs were counted after three days.

### 2.3. DNA extraction, sequencing, and construction of reference genomes

To obtain draft genomes for RNA read mapping, 5 mL *A. baumannii* or *K. pneumoniae* and 15 mL *Bacillus* spp. DNA was extracted using the DNeasy PowerSoil Pro Kit (Qiagen) following the manufacturer’s instructions. DNA samples were submitted to the Microbial Genome Sequencing Center (MiGS) for library construction and short-read sequencing on an Illumina platform. KneadData v0.7.10 (https://bitbucket.org/biobakery/kneaddata/) was then used for raw sequence quality control and contaminant removal with default parameters. Short reads were assembled into contigs using SPAdes v3.14.1^36^ and each reference genome was annotated using Prokka v1.14.6^37^. Additional annotation of antibiotic and antimicrobial resistance genes were conducted using the Comprehensive Antibiotic Resistance Database (CARD)^38^. Assembled contigs from All Purpose Probiotic Cleaner were also binned into five metagenome-assembled genomes (MAGs) using metaWRAP^39^. CheckM^40^ was used to assess the quality of microbial genomes. Taxonomic classification of each draft genome was determined by GTDB-Tk^41^.

### 2.4. RNA Extraction, Sequencing, and Data Processing

Swab samples were stored in RNAprotect Bacteria Reagent under -80°C until extraction. RNA was extracted using the RNeasy Mini Kit (Qiagen) following its recommended “Protocol 2: Enzymatic lysis and mechanical disruption of bacteria’’, consisting of a 10 min treatment with 15 mg/mL lysozyme and 10 min beadbeating. Libraries were constructed at Northwestern NUSeq using the Illumina Total RNA Prep with Ribo-Zero Plus, according to manufacturers’ protocol. A total of 110 samples (108 samples collected at various time points and cleaning scenarios, one kit control, and one sample prepared for between-batch normalization) were sequenced using an Illumina HiSeq4000 with 50 bp single end reads.

Reference genomes constructed from whole genome sequencing were used for RNA mapping. Raw sequence files were processed using trim_galore v0.6.6 (https://www.bioinformatics.babraham.ac.uk/projects/trim_galore/) and Kneaddata v0.7.10 (https://bitbucket.org/biobakery/kneaddata/) to remove sequence adapters, low quality reads, and contaminants. Sortmerna v4.2.0^42^ was used to filter rRNA reads against the silva-bac-16s-id90 and silva-bac-23s-id98 databases. HISAT2 v2.1.0^43^ was used to build local indices and map short reads onto reference draft genomes. Transcript abundances were quantified using featureCounts v2.0.1^44^. Differentially expressed genes (DEGs) were identified by DESeq2^45^ at a significance level of 0.05 (Benjamini-Hochberg adjusted p-value). BlastKOALA (KEGG Orthology And Links Annotation) was used for K number assignment followed by Prokka annotation^46^. KEGG pathway enrichment analysis was further conducted using ClusterProfiler^47^. Statistical analysis, including Welch’s paired two sample *t*-test, principal component analysis (PCA), and permutational multivariate analysis of variance (PERMANOVA, 9,999 permutations) were conducted in R (version 4.1.2, 2021-11-01).

## 3. Results and Discussion

### 3.1. Survival of *A. baumannii* and *K. pneumoniae* under four cleaning scenarios and two temperature/humidity conditions

We quantified the CFUs of *A. baumannii* and *K. pneumoniae* to examine their survival under ambient and humid conditions in combination with four cleaning strategies as listed in Table 1. Over the course of 3 days, we observed a maximum log_10_ reduction of 8.75 for *A. baumannii* (Cleaner Only and Probiotic Cleaner under ambient condition) and 7.42 for *K. pneumoniae* (No Cleaner, Cleaner Only and Probiotic Cleaner groups). Under ambient conditions, both Cleaner Only and Probiotic Cleaner demonstrated a near-complete inactivation of *A. baumannii* within 72 hours (<LOD) and complete inactivation for *K. pneumoniae*. Although the survival of *A. baumannii* between Probiotic Cleaner and Cleaner Only are not statistically significant (p>0.05, Figure S1), Probiotic Cleaner cleaning at 24 hours showed 1.43 greater log_10_ reduction compared to Cleaner Only (Figure 2A). As there was no significant difference between probiotic cleaning and cleaner only, the reduction of *A. baumannii* was largely attributed to the chemical component of the cleaning product. Differences in the survival of *K. pneumoniae* between Probiotic Cleaner, Cleaner Only and No Cleaning scenarios was not statistically significant (p>0.05, Figure S1), with only 0.77 greater log_10_ reduction in Probiotic Cleaner compared to Cleaner Only at 24 hours. The fact that *K. pneumoniae* was unable to persist on the surface in No Cleaner group under ambient conditions indicated that chemical components in the cleaning product only had marginal contribution to *K. pneumoniae* inactivation.

**Figure 2.**
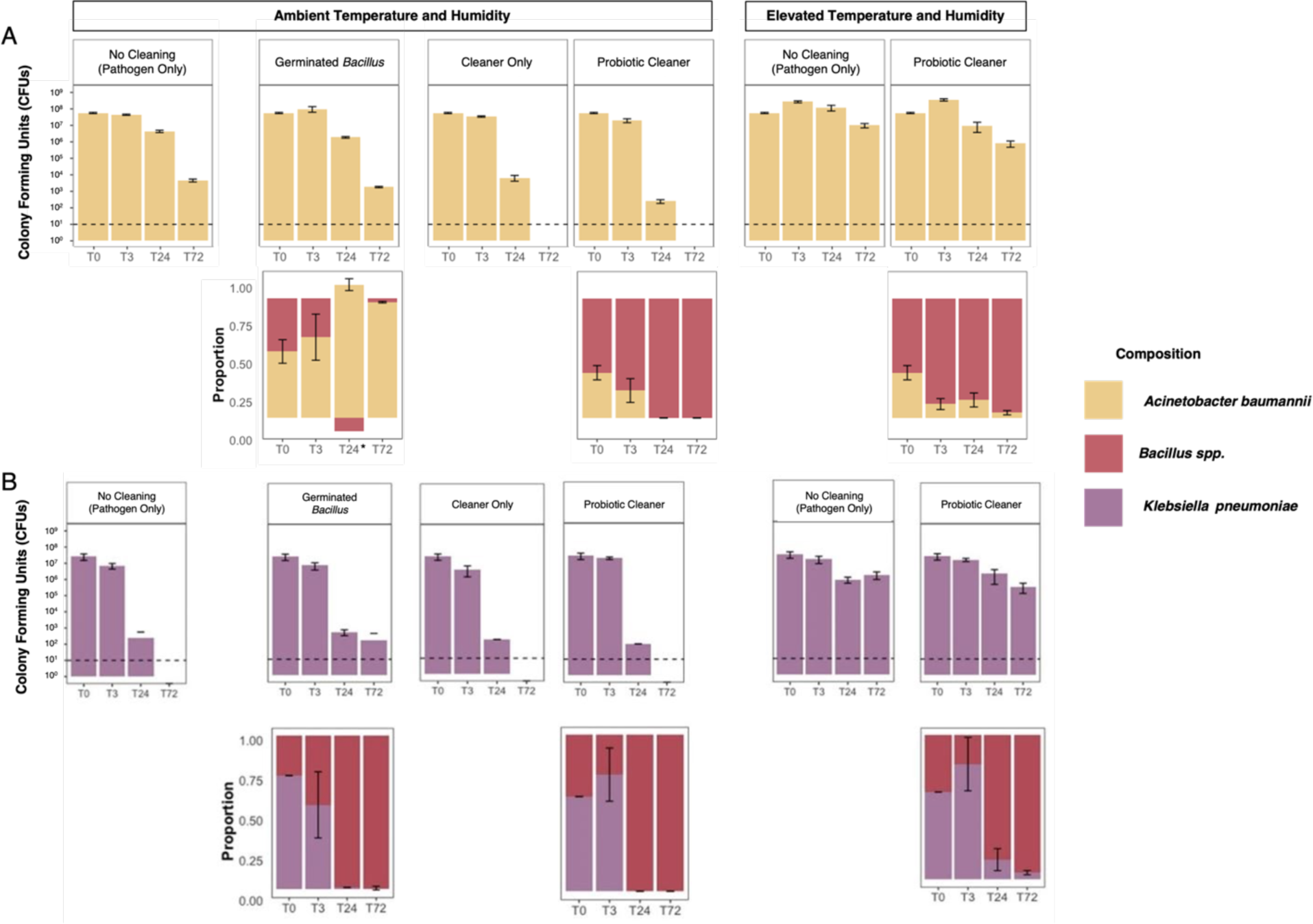
Colony forming units (CFU) of pathogens and composition of surface microbial community under four cleaning scenarios and two temperature and humidity conditions. CFUs of pathogens were counted on day 3 of incubation (37°C) on their selective media while total CFUs (pathogen and *Bacillus* spp.) were obtained from TSA. The proportions of *A. baumannii* and *K. pneumoiae* were calculated as the ratio between CFU from their selective media and CFU from TSA. The proportion of *Bacillus* spp. was calculated as 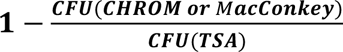 . Dashed lines represent a limit of detection (LOD) of 10 CFU. * Due to possible technical variation, the proportion of *A. baumannii* exceeded 1 at T24 for *A. baumannii* Germinated *Bacillus* scenario.

Surfaces cleaned with Probiotic Cleaner were also gradually dominated by *Bacillus* spp. due to the decreasing absolute abundance of the pathogen population. This persistence of *Bacillus* spp. on surfaces was observed under both ambient and elevated humidity and temperature (Figure 2). In contrast, the germinated *Bacillus* spp. spores introduced as vegetative cells were gradually taken over by the *A. baumannii* population; for *K. pneumoniae* Germinated Probiotic group the cleaned surface was still dominated by *Bacillus* spp. over *K. pneumoniae* (Figure 2A). Without the use of chemical-based surface cleaner (No Cleaner and Germinated *Bacillus*), *A. baumannii* showed comparable CFUs at all time points (p > 0.05).

Throughout the experiments, the minimum log_10_ reduction was observed under elevated temperature and humidity for both *A. baumannii* and *K. pneumoniae*. Their survival over time improved under elevated temperature and humidity with or without probiotic cleaner (Figure 2) although *A. baumannii* exhibited higher viable populations (p<0.01, Figure S1A) while *K. pneumoniae* showed low or no significance (Figure S1B) when comparing No Cleaner (Humid) to No Cleaner (Ambient) (paired *t*-test was performed for all time points). Higher viability of bacteria is observed under higher RH, which is related to the stress caused by concentrating of solute and desiccation as droplets dry out^48^. Genes *csrA* which is related to carbon storage and *bfmR* (*rstA*) which is related to stress response play a major role in desiccation tolerance in *A. baumannii* ^49, 50^. Both genes were downregulated under humid condition in *A. baumannii* No Cleaner groups compared to ambient condition, and *csrA* gene was also downregulated under humid condition in *K. pneumoniae* Probiotic Cleaner group compared to ambient condition (log_2_ fold change value < -1, adjusted p-value < 0.05, data not shown). This result suggests greater potential health risks and infectious disease burdens in developing regions with a humid and hot climate and limited climate control^51^.

### 3.2. Overview of whole genome sequencing and metatranscriptomic data

Sequencing of the *A. baumannii* isolate generated 3.72 million reads with an average read length of 136 bp. Reads were assembled into contigs with a N50 of 141,238 bp. Sequencing of *K. pneumoniae* generated 4.4 million reads with an average rate of 145 bp. Reads were assembled into contigs with a N50 of 113,698 bp. Sequencing of the All-Purpose Probiotic Cleaner resulted in 16.49 million reads with an average read length of 139 bp, after quality trimming and contaminant removal. Reads were assembled into contigs with a N50 of 143,313 bp. Open reading frame (ORF) identification recognized 4,039, 5,823, and 22,429 coding regions for *A. baumannii, K. pneumoniae,* and probiotic cleaner, respectively, 2,253, 4052, and 13,339 of which were functionally annotated (not as “hypothetical protein”). Among these annotated regions for *A. baumannii, K. pneumoniae,* and probiotic cleaner assemblies, 2,150, 3,675, and 11,594 were classified into one or more KEGG Orthologs respectively. Contigs from probiotic cleaner reads were binned into five groups with completeness > 90%, and contamination < 5%. All five bins were classified as *Bacillus* species as shown in Table 2.

**Table 2.** *Bacillus* genome binning and taxonomic assignment. Completeness and contamination statistics were determined using the CheckM algorithm.

| Bin | Completeness<br>(%) | Contamination<br>(%) | GC<br>(%) | Size<br>(Mbp) | Taxonomic Classification |
| --- | --- | --- | --- | --- | --- |
| 1 | 90.58 | 1.883 | 0.351 | 4.74 | *Bacillus cereus* |
| **2** | 96.43 | 1.489 | 0.466 | 3.60 | *Bacillus velezensis* |
| **3** | 99.11 | 0.148 | 0.436 | 4.11 | *Bacillus subtilis* |
| **4** | 98.21 | 0.087 | 0.461 | 4.28 | *Bacillus licheniformis* |
| **5** | 99.08 | 0.207 | 0.416 | 3.57 | *Bacillus safensis* |

RNA samples containing *A. baumannii* or *K. pneumoniae* were collected for up to 72 hours of surface inoculation. Biological triplicates were collected for each condition and an average of 12.73 ± 3.83 and 11.03 ± 4.98 million reads with an average read length of 49 bp were recovered for *A. baumannii* and *K. pneumoniae* experiments, respectively, after removing low-quality reads, contaminated sequences, adapters, and rRNA. The average mapping rate using HISAT2 was 98.20% ± 1.06% for experiments with *A. baumannii* and 96.40% ± 10.24% for experiments with *K. pneumoniae* (Figure S2). Percentages of reads mapped onto *A. baumannii* and *Bacillus* genomes vary across four cleaning scenarios; Germinated *Bacillus* samples containing vegetative *Bacillus* spp. on average had 40.53% ± 6.07% reads mapped onto *Bacillus* reference genomes, whereas this number for Probiotic Cleaner samples was only 0.56% ± 0.75%. A similar trend was observed in experiments with *K. pneumoniae*. Samples containing *K. pneumoniae* and vegetative *Bacillus* spp. on average had 40.52% ±16.83% reads mapped onto *Bacillus* reference genomes; however, this number was only 1.00 ±1.08% for Probiotic Cleaner samples. Thus, these data suggest that the majority of *Bacillus* spp. in the probiotic product likely remained as spores throughout the incubation. Due to this insufficient coverage of *Bacillus* transcriptome in Probiotic Cleaner samples, our transcriptomic analysis is composed of quantitative comparisons for the *A. baumannii* and *K. pneumoniae* transcriptomes and qualitative evaluations for *Bacillus* spp.

### 3.3. Transcriptomes of *A. baumannii* and *K. pneumoniae* clustered into distinct groups determined by cleaning and physical conditions

Prior to conducting principal component analysis (PCA), read count data were transformed to remove the dependence of the variance on the mean. PCA of the transformed (variance stabilization) read counts belonging to *A. baumannii* indicated that the expression of *A. baumannii* clustered into three distinct groups based on cleaning scenarios and physical conditions (Figure 3A). Clustering of read counts from *K. pneumoniae* is more distinct for different time points and physical conditions (Figure S2) compared to cleaning scenarios and physical conditions (Figure 3C). PERMANOVA (9,999 permutations) revealed that temperature and humidity (p<0.001), cleaning (p<0.001), and time (p<0.001) explained 12%, 30%, and 9% of the variance for *A. baumannii* read counts, and 29%, 10%, and 12% for *K. pneumoniae* read counts, respectively. Gene expression profiles of *A. baumannii* with detergent cleaning (Probiotic Cleaner and Cleaner Only) are more similar to each other than they were to those without detergent (Germinated *Bacillus*). Temperature and humidity conditions also affected the expression of *A. baumannii* under the no-cleaning condition, but this impact was minimal when probiotic cleaner was used. On the contrary, the temperature and humidity strongly affected the gene expression profiles of *K. pneumoniae*, and the impact was more pronounced as the cleaning time extended from 3 hours to 24 and 72 hours.

**Figure 3.**
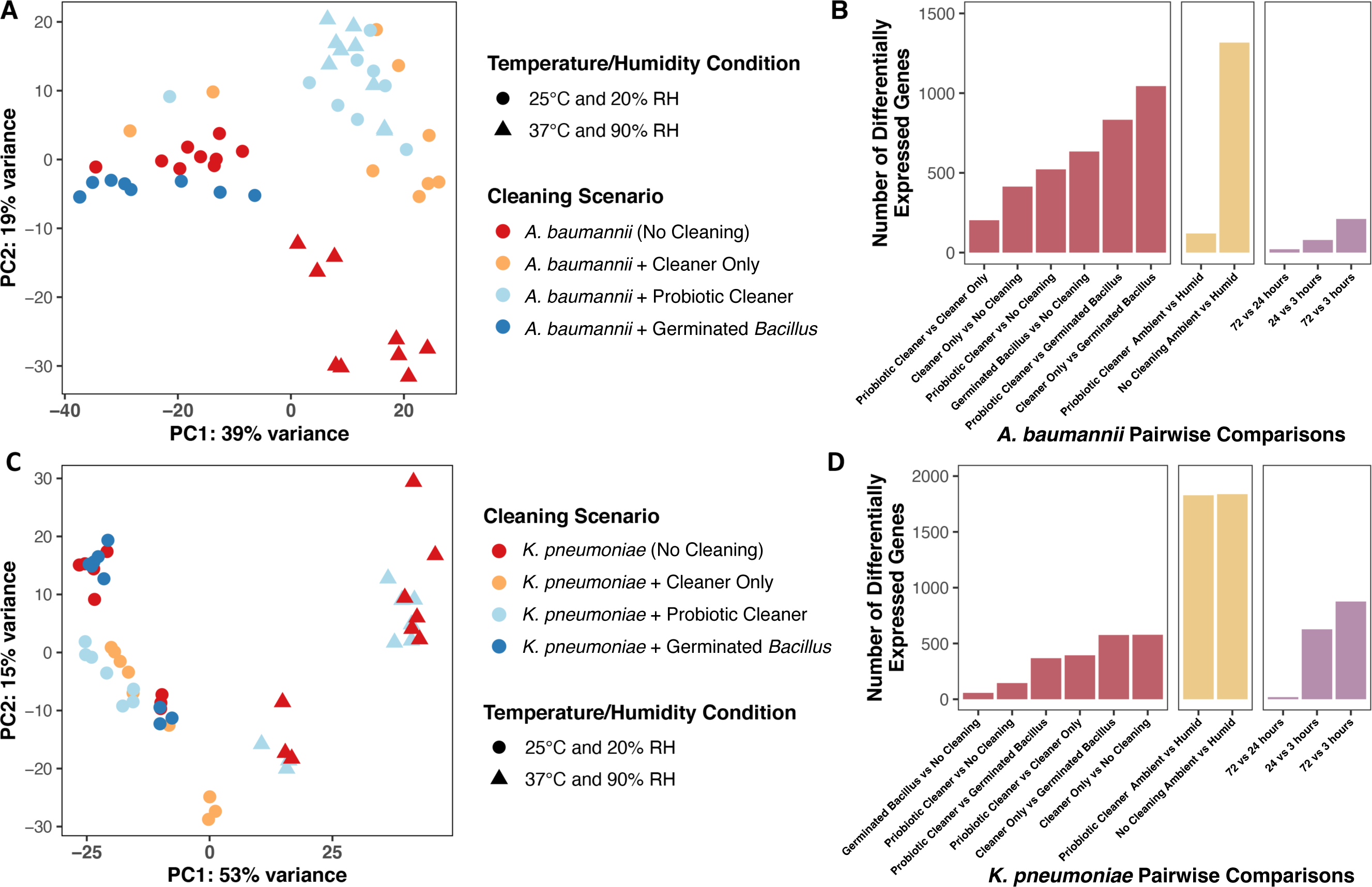
Principal component analysis on variance stabilized *A. baumannii* (A) and *K. pneumoniae* (C) read counts and number of differentially expressed genes of 11 pairwise comparisons for *A. baumannii* (B) and *K. pneumoniae* (D) experiments. Samples were color-coded based on cleaning conditions; each point (dot or triangle) represents its corresponding temperature and humidity condition.

Using No Cleaner samples as the reference transcriptome, genes with absolute log_2_ fold change values greater than or equal to 1 as calculated by DESeq2 were considered as differentially expressed genes (DEGs, Benjamini-Hochberg adjusted p-value ≤ 0.05). 414, 522, and 634 genes of *A. baumannii* and 578, 145, and 57 genes of *K. pneumoniae* were found to be differentially expressed in Cleaner Only, Probiotic Cleaner, and Germinated *Bacillus* samples, respectively. The transcriptome of *A. baumannii* treated by probiotic cleaner and cleaner only shared the highest level of similarity and consequently, the smallest number of differentially expressed genes (Probiotic Cleaner *vs* Cleaner Only, Figure 3B). In the case of *K. pneumoniae*, Germinated *Bacillus* samples were the most similar to No Cleaner samples. Consistent with previous PCA analysis, expression of *A. baumannii* under ambient and elevated temperature and humidity conditions are drastically different from each other (1298 DEGs, Figure 3B) whereas this difference in transcriptome diminished when probiotic cleaner was used (133 DEGs). The high impact of elevated temperature and humidity persisted for *K. pneumoniae* even when probiotic cleaner was applied (1828 and 1838 DEGs with or without probiotic cleaner, Figure 3D). Since the transcriptome of *K. pneumoniae* under Probiotic Cleaner scenario was fairly similar to the No Cleaner scenario, their similar response to elevated temperature and humidity could be expected. Samples collected at various time points throughout the microcosm inoculation also showed that much of the temporal variation in gene expression occurred within 24 hours of surface cleaning for *A. baumannii* (Figure 3B), *K. pneumoniae* (Figure 3D), and vegetative *Bacillus* (Figure S4), as the number of DEGs of the 72-hour *vs*. 24-hour comparison is much lower compared to 24- hour *vs*. 3-hour and 72-hour *vs*. 3-hour comparisons. During the experiment, we typically observed drying out of the bacterial liquid culture droplets within 24 hours. This result is consistent with study by Lin and Marr^52^ on bacteria viability where bacteria inactivation rate is higher in evaporating droplets compared to when droplets are dried out. Based on the observation, cleaning protocols and environmental conditions should be considered for effective disinfection in addition to cleaning reagent types.

### 3.4. KEGG pathway enrichment profiles revealed different metabolic response under different cleaning scenarios

Enriched transcriptomic signatures varied greatly across cleaning conditions and between the tested pathogens, *A. baumannii* and *K. pneumoniae*. Overall, the *K. pneumoniae* transcriptome responded with greater heterogeneity to treatment than the *A. baumannii* transcriptome, as evidenced by their distinct patterns of treatment-specific enrichment (Fig. 4A and 4D top portion; e.g., *A. baumannii* with Germinated Probiotic). These phenomena support the hypothesis that the *A. baumannii* genome encodes survival mechanisms for persistence in replete environments, while *K. pneumoniae*, although capable of short-term response, may be less equipped for long term survival on nutrient poor surfaces.

**Figure 4.**
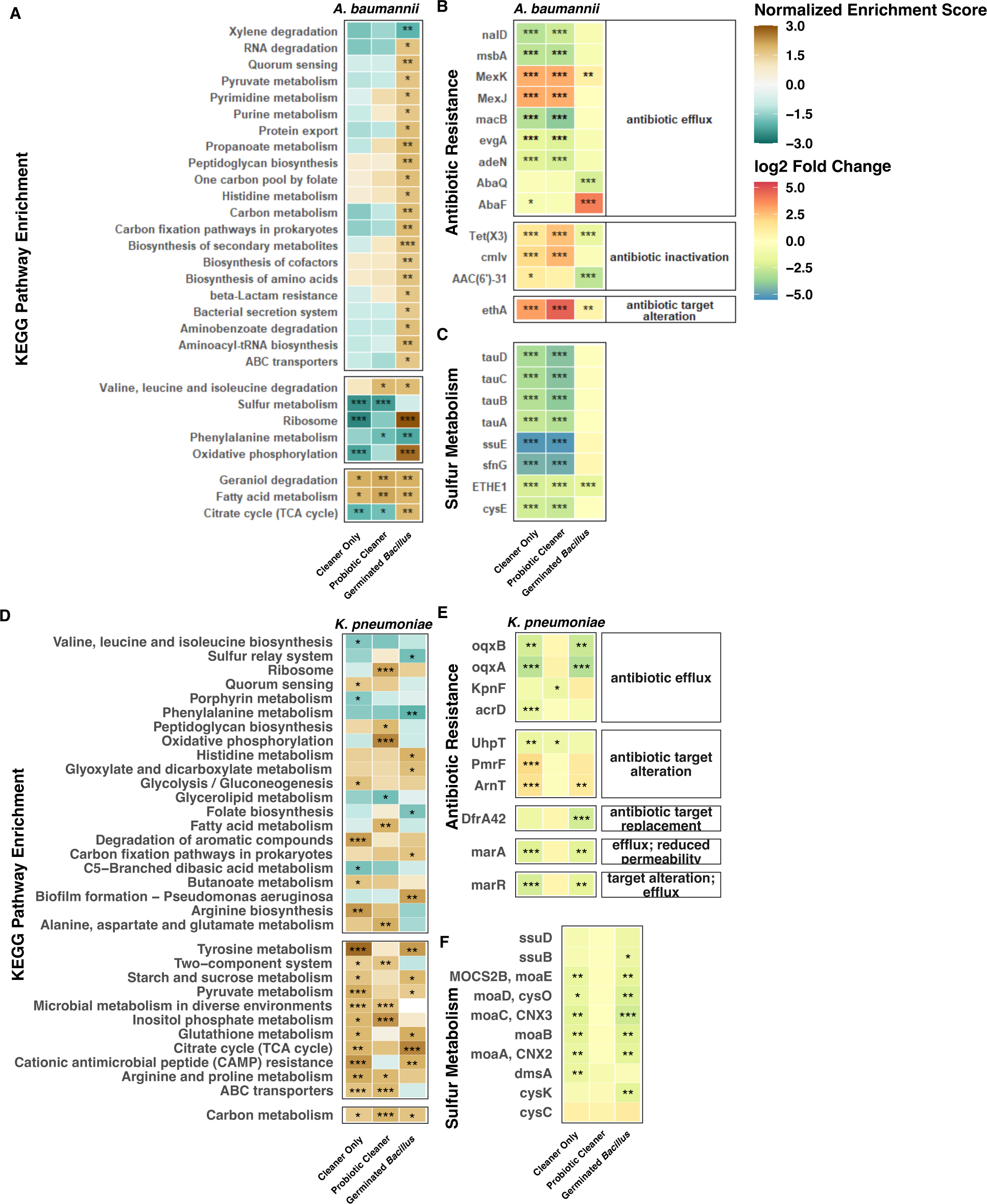
KEGG pathway enrichment results for samples collected under ambient temperature and humidity condition (A, D), log_2_ fold change of genes associated with antibiotic/antimicrobial resistance (B, E), and sulfur metabolism (C, F), compared to no-cleaning samples (*A. baumannii* or *K. pneumoniae*). KEGG pathways shown here were statistically enriched in at least one of the cleaning conditions (Benjamini-Hochberg corrected p-value ≤ 0.05). Normalized enrichment score (NES) accounts for differences in gene set size and were used to compare enrichment results across KEGG pathways; a positive NES suggests an overall positive upregulation of genes belonging to its corresponding pathway^47, 70, 71^. * p<0.05, ** p< 0.01, and ***p < 0.001.

The *A. baumannii* transcriptome shifted under treatment conditions and 26 KEGG pathways broadly related to metabolism, energy production, translation, and membrane transport were enriched. Significant enrichment of translation machinery under Germinated Probiotic treatment, suggests that protein synthesis in *A. baumannii* was likely promoted to achieve a large demand for surface competitions. On the other hand, three pathways were positively enriched in samples with chemical-based detergent cleaning, indicating that the *A. baumannii* population overall repressed metabolic activities and adaptively reduced synthesis in response to environment constraints associated with detergent cleaning (Figure 4A). As *A. baumannii* is notoriously associated with the hospital environment^53^, it stands to reason that it may remain constitutively competent to respond to such replete environments and have acquired additional adaptive traits to quickly acclimate when additional stressors, e.g., chemical cleaners, common in the hospital environment arise.

In contrast, *K. pneumoniae* responded with far greater heterogeneity at the transcript level. Carbon metabolism genes in *K. pneumoniae* were mutually upregulated in every cleaning condition; however, specific carbon metabolism pathways were induced in a treatment-dependent manner. For instance, TCA cycle, pyruvate, starch and sucrose metabolism genes were upregulated by treatment with germinated *Bacillus* or cleaner only but not in samples treated with probiotic cleaner. This suggests that vegetative *Bacillus* and cleaning products induce alternative though overlapping carbon metabolism networks in *K. pneumoniae*. Indeed, increased expression of biosynthetic pathways (glyoxylate cycle, carbon fixation) suggests vegetative *Bacillus* may promote anabolic competency in *K. pneumoniae.* Perhaps this broad range of metabolic activity is the result of slight alterations in the *K. pneumoniae* panic response, as the population tries to survive in a “hostile” environment, unlike that of its native host.

While fewer pathways (11) were uniquely enriched in response to germinated *Bacillus,* every treatment stimulated some unique pathway upregulation in *K. pneumoniae,* unlike the uniform pattern observed in *A. baumannii* (Fig 4D). Genes involved in two-component systems and membrane transport (ABC) were upregulated in samples treated with probiotic cleaner or cleaner only but not in samples containing vegetative *Bacillus*, indicating that chemical detergents, rather than *Bacillus*, induce environmental sensing and response mechanisms in *K. pneumoniae. Surprisingly*, no pathways were mutually and specifically enriched by vegetative *Bacillus* or probiotic cleaner, which may be influenced by the diminished transcriptional landscape of *Bacillus* spores (Figure S2). Alternatively, the divergent transcriptomic profiles induced by each cleaning condition indicates that *K. pneumoniae*’s strategy for surviving on surface environments vastly differs from *A. baumanii* and further suggests that the *A. baumanii* response system to built environment surfaces may be more robust than *K. pneumoniae*’s. Indeed, evolutionary studies modelling extended resource exhaustion in *Escherichia coli* have shown that while stress responses dominate in the short-term, in the long-term, genetically-encoded adaptations to replete environments sweep the population via clonal expansion^54, 55^. One could thus hypothesize the dissimilar fitness levels and phenotypic responses (e.g., enrichment patterns) of *K. pneumoniae*^56^ and *A. baumanii* in this study may be the results of their unique evolutionary history and ecological niches.

Cleaning products may represent a source of nutrients in an otherwise replete environment. Both *A. baumannii* and *K. pneumoniae* exhibited increased transcription of genes related to metabolism, although not the same pathways and not under all conditions. Our results suggest that vegetative *Bacillus* survive poorly on surfaces and become a source of nutrients to more resilient *A. baumannii. K. pneumoniae* enhances transcription of various metabolic pathways, but none of the responses enhance its survival. The specific metabolic activity of organisms on surfaces is highly dependent on both the organism and the resource availability.

### 3.5. Different cleaning scenarios induced differential expression of genes associated with stress response and competition

We examined gene expression of important interspecies competition pathways to understand the potential interactions between probiotic bacteria *Bacillus* spp. and pathogens *A. baumannii* or *K. pneumoniae*. The two pathogens responded differently toward cleaning scenarios where *A. baumannii* exhibited more pronounced differential gene expression compared to *K. pneumoniae*.

Iron acquisition through siderophores is believed to be one of the key mechanisms of inter-species competitive exclusion^57^. Differential expression of the siderophore and iron acquisition genes indicated that siderophore cheating is unlikely to happen under our studied conditions. In *A. baumannii* experiments, several genes responsible for siderophore synthesis, export, and reception (e.g., aerobactin, anguibactin, and enterobactin) were downregulated in the Germinated Probiotic group (Figure S5A). Although bacteria with multiple siderophore receptors can gain competitive advantages in social competition through siderophore cheating^58^, the downregulation of siderophore-related genes in these samples was likely due to utilization of iron originating from lysed cell debris, especially from *Bacillus* spp., rather than active competition through siderophore cheating, given the inability of vegetative *Bacillus* spp. to persist on surfaces (Figure 2A). For *K. pneumoniae* experiments, no siderophore-related gene was differentially expressed under the three cleaning scenarios (Figure S6A, *iutA*, *fepA*, and *fepC* log_2_ fold change absolute values < 1). Gene expression levels of *Bacillus* spp. (Figure S7) further precluded siderophore cheating as the competing mechanism on surface as genes related to the synthesis, and export of *Acinetobacter-* and *Klebsiella*-associated siderophores (e.g., aerobactin, anguibactin, and enterobactin) were only occasionally expressed (Figure S7).

In addition to iron acquisition, genes related to Type VI Secretion Systems (T6SS), Type IV or Type 1 Pili, and biofilm formation were shown to have important implications in inter-species competition, virulence expression, and starvation response in *A. baumannii* or *K. pneumoniae* ^57, 59, 60^. In *A. baumannii* experiments, genes associated with T6SS, Type IV Pili, poly-beta-1,6-N-acetyl-D-glucosamine (PGA) production and biofim formation were upregulated in Probiotic Cleaner, Cleaner Only, and Germinated Probiotic samples with varying statistical significance (Figure S5). The *pgaABCD* operons, which had increased expression in Germinated Probiotic group, play roles in surface binding and maintaining biofilm structure stability^61, 62^. This result suggested greater potential for PGA, a biofilm adhesin polysaccharide^61, 62^, production and subsequent biofilm activities when *A. baumannii* and *Bacillus* spp. co-existed. In contrast, *K. pneumoniae* exhibited negligible differences in expression levels for genes related to T6SS and Type 1 Pili had under different cleaning scenarios for (Figure S6BD). Additionally, only a few biofilm formation related genes were differentially expressed (Figure S7C). Lack of shared differentially expressed genes across cleaning scenarios indicated nonspecific response of *K. pneumoniae* biofilm formation towards different stressors.

Similar to what was observed for KEGG pathways, antibiotic resistance gene expression of probiotic cleaner and cleaner only treated *A. baumannii* exhibited high similarity (Figure 4B) while *K. pneumoniae* responded very differently to probiotic cleaner and clear only treatments (Figure 4E) in. In the Germinated Probiotic groups, we observed an increased expression of *abaF*, a gene encoding fosfomycin resistance, in *A. baumannii* while no antibiotic resistance gene was upregulated (log_2_ fold change ≥ 1) in *K. pneumoniae*. Interestingly, no genetic determinants of fosfomycin biosynthesis were identified from *Bacillus* spp. reference genomes. In addition, it was reported that the disruption of *abaF* in *A. baumannii* not only resulted in an increase in fosfomycin susceptibility but also a decrease in biofilm formation and virulence^63^. Therefore, the upregulation of *abaF* in *A. baumannii* may not be induced by the fosfomycin secretion from *Bacillus* spp. but rather a stress response to *Bacillus* spp. existence exerting enhanced biofilm formation.

In *A. baumannii* experiments, pronounced downregulation of genes related to sulfur metabolism, particularly those induced by sulfur starvation, was observed for groups where chemical cleaning detergent was present (Probiotic Cleaner and Cleaner Only groups for the ambient condition, Figure 4C; Probiotic Cleaner group for the humid condition, Figure 5C). These genes include *ssuEADCB* and *tauABCD* operons known to assimilate sulfur from aliphatic sulfonates^64^ and facilitate the utilization of taurine as an alternative sulfur source under sulfate-deprived environment^65^. Owing to the presence of anionic surfactants in liquid detergent products, such as sodium lauryl ether sulfate (SLES)^66^, these sulfate starvation genes were likely suppressed in Cleaner Only and Probiotic Cleaner groups. This observation is consistent with studies that illustrated the overexpression of *tauABCD* and *ssuEADCB* genes induced by sulfate shortage^67^. Additionally, bacterial utilization of SLES facilitated by *Citrobacter braakii* ^68^ and a consortium of *Acinetobacter calcoacetiacus*, *Klebsiella oxytoca*, and *Serratia odorifera*^66^ has also been reported in previous studies and may contribute to the persistence of microbial communities associated with hospital sinks^69^. On the contrary, no differential expression was observed for sulfur metabolism genes in *K. pneumoniae* Probiotic Cleaner group under both ambient and humid conditions (Figure 4F and Figure 5F).

**Figure 5.**
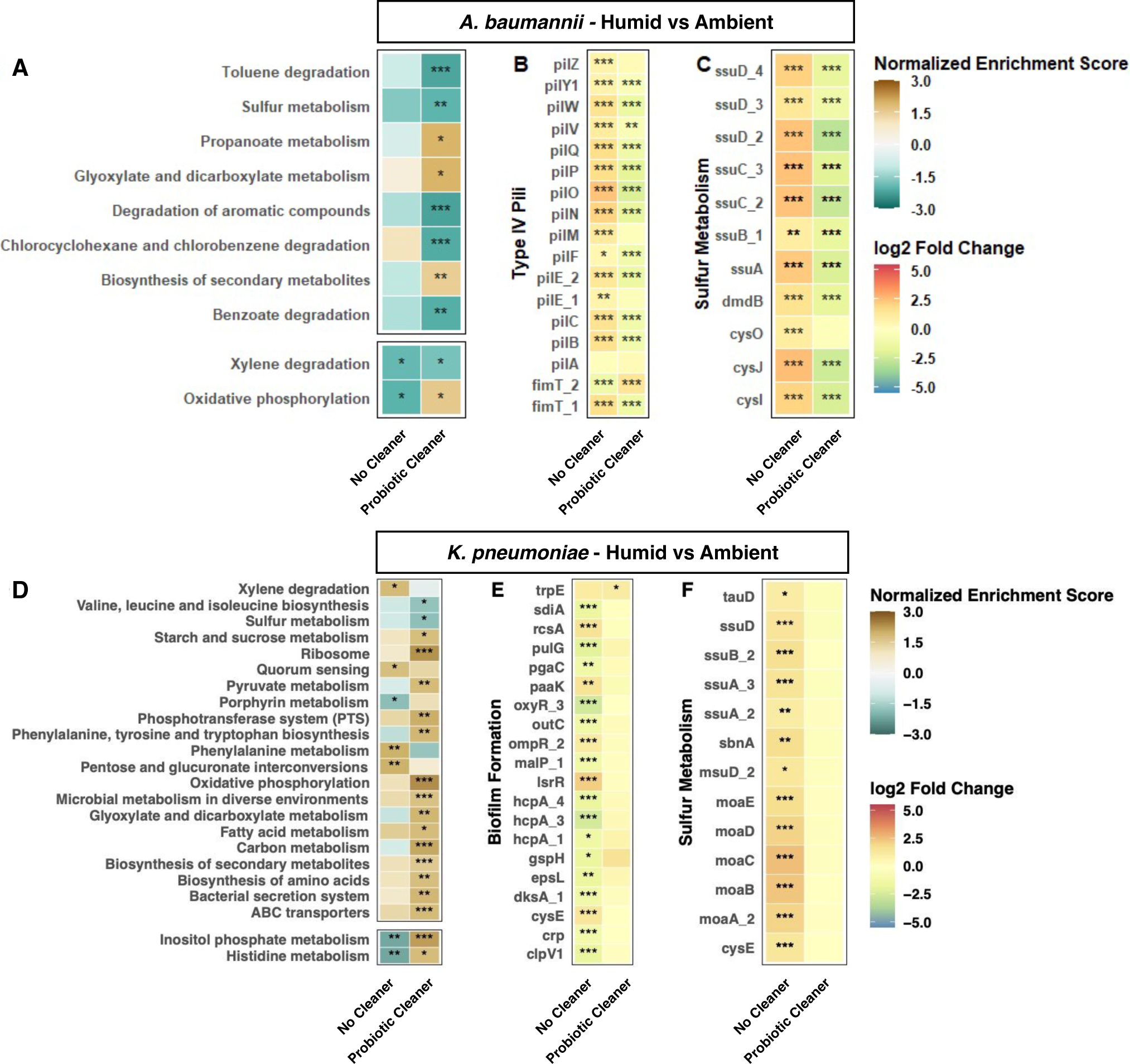
KEGG pathway enrichment results for No Cleaner and Probiotic Cleaner samples collected under elevated temperature and humidity condition (A, D), log_2_ fold change of genes associated with *A. baumannii* cell motility (B), *K. pneumonia*e biofilm formation (E), and sulfur metabolism (C, F). KEGG pathways shown here were statistically enriched in at least one of the cleaning conditions (Benjamini-Hochberg corrected p-value ≤ 0.05). * p<0.05, ** p< 0.01, ***p < 0.001.

In *K. pneumoniae* experiments, Cleaner Only and Germinated Probiotic groups showed similarities in gene expressions while Probiotic Cleaner appeared to have very limited effects on *K. pneumoniae* gene expression. It is possible that the dormant *Bacillus* spores in probiotic cleaner were buffering the stress caused by chemical cleaner while the vegetative *Bacillus* cells introduced competition pressure on the surfaces. Thus, it is important to understand how spores and vegetative probiotic microorganisms interact with and affect the performance of chemical surface cleaning reagents, which can alter the response landscape of the surface associated pathogens. Since the results from our study showed species-specific responses, more pathogen species should be investigated for their interspecies interactions with probiotic microorganisms to observe general patterns. This will set the foundation and facilitate the development of targeted cleaning and disinfection strategies.

### 3.6. Effects of elevated temperature and humidity on pathogen cell motility and biofilm formation

Although over 1,000 differentially expressed genes were identified from *A. baumannii* or *K. pneumoniae* transcriptomes between the ambient and humid conditions, the majority of the KEGG pathways were composed of genes that were both upregulated and downregulated, rendering non-significant results for most pathways.

Under the humid condition (elevated temperature and humidity), 16 KEGG pathways related to metabolism (10), membrane transport (3), biosynthesis (2) and translation (1) were enriched in *K. pneumoniae* experiments (Figure 5D, Probiotic Cleaner Humid *vs* No Cleaner Humid) while only 4 KEGG pathways related to metabolism (2), biosynthesis (1) and energy production (1) were enriched in *A. baumannii* (Figure 5A, Probiotic Cleaner Humid *vs A. baumannii* Humid) with Probiotic Cleaner. Overall, more KEGG pathways were affected by elevated temperature and humidity in *K. pneumoniae* than *A. baumannii* (Figure 5D and Figure 5A). This result corresponded to the PCA and PERMANOVA results where temperature and humidity affected the transcriptome profiles of *K. pneumoniae* more than *A. baumannii*, but not reflected in the survival of the two pathogens under elevated temperature and humidity assessed by viable CFU counts.

Without Probiotic Cleaner, cell motility appeared to be affected differently by increased moisture availability for *A. baumannii* and *K. pneumoniae*. *A. baumannii* exhibited upregulation of type IV pili genes (Figure 5B) whereas *K. pneumoniae* type 1 pili genes remained unaffected, which was also the case under ambient condition (Figure S6D). Nevertheless, 19 genes related to biofilm formation were differentially expressed (Figure 5E, *K. pneumoniae* Humid vs *K. pneumoniae* Ambient), and the quorum sensing pathway, which affects biofilm formation of *K. pneumoniae*^72^ , was enriched under elevated temperature and humidity (Figure 5D, *K. pneumoniae* Humid vs *K. pneumoniae* Ambient). This indicated that elevated temperature and humidity could assist the persistence of *K. pneumoniae* on surfaces (Figure 2B) by altering the biofilm formation related gene expression.

For both *A. baumannii* and *K. pneumoniae* under Probiotic Cleaner scenario, sulfur metabolism pathway was downregulated under elevated temperature and humidity, which is consistent with the sulfur metabolism-related gene expression results discussed in the previous section when chemical detergent was present. The effects of probiotic cleaner on *K. pneumoniae* were likely masked by the changes due to higher temperature and humidity (Figure 5E and Figure 5F), meaning the benefits of probiotic cleaning products could be more limited in tropical areas for certain types of pathogens.

Our results demonstrated the persistence of probiotic *Bacillus* included in the All-Purpose Probiotic Cleaner up to 72 hours after cleaning. However, this persistence did not result in more efficient reduction of pathogen biomass. *A. baumannii* on surfaces cleaned using chemical-based detergent with and without probiotic *Bacillus* contained a comparable amount of viable pathogens. The transcriptome of *A. baumannii* with and without probiotic addition shared a high degree of similarity in overall gene expression. In contrast, the transcriptome of *K. pneumoniae* with probiotic addition showed a high degree of differences compared to samples with chemical cleaner only or germinated *Bacillus*, including reduced response in genes related to antimicrobial resistance, sulfur metabolism and biofilm formation. We thus conclude *Bacillus* spores indeed survive unfavorable conditions, but that the small percentage of germinated *Bacillus* cells and/or slow germination rate on surfaces preclude interspecies interactions like competitive exclusion. Elevated temperature and humidity resulted in prolonged persistence of viable pathogens on surfaces. These conditions also had a pronounced effect on *K. pneumoniae* overall gene expression regardless of which cleaner was used. These results suggest that vigilant indoor climate control could contribute to infection prevention. Especially in tropical environments, our results suggest that reducing the temperature and relative humidity is a more important strategy for infection prevention than cleaning product selection.

The design of more effective probiotic cleaning products and strategies to promote “healthy” microorganisms will require a more nuanced understanding of the relevant environmental conditions and how they impact activity. Our study used microcosms to model environmental conditions *in situ* and enable the manipulation of individual parameters. While our microcosms do not represent the life histories of organisms *in situ* (e.g., previous exposure and selection on a surface), this type of work forms an important bridge between experiments in culture media, which do not reflect *in situ* conditions, and field studies, which preclude replication and manipulation. We further demonstrate that transcriptomics can be used to assess activity on surfaces. We advocate for the inclusion of such assessments in field studies, which are historically typically DNA- or culture-based, especially as RNA sequencing methods improve for low biomass samples. The assessment of cleaning and disinfection is currently based on log reductions of viable biomass. Using these tools to progress beyond taxonomic and viability profiling will allow us to include more nuanced endpoints, such as the prevention of difficult-to-disinfect phenotypes like biofilm formation or antimicrobial resistance.

## Acknowledgement

This work was supported, in whole or in part, by the Bill & Melinda Gates Foundation INV-004298. Under the grant conditions of the Foundation, a Creative Commons Attribution 4.0 Generic License has already been assigned to the Author Accepted Manuscript version that might arise from this submission. JH is supported in part by a Terminal Year Fellowship from the Northwestern University McCormick School of Engineering. The authors would like to thank Marina Zelivyanskaya and Dr. Alan R. Hauser for providing clinical isolates of *Acinetobacter baumannii* and *Klebsiella pneumoniae*. We further thank Andreas Wagner and Jiaxian Shen for their assistance.

## Author Contributions

JH designed and performed the experiments, analyzed data, and wrote the manuscript. WS, JTS, and AAM analyzed transcriptomics data and wrote the manuscript. EMH supervised the project.

## Data Availability

Raw sequencing data are available in NCBI Sequence Read Archive (SRA) under BioProject accession number PRJNA818256. Processed data files and scripts for data analysis are available in GitHub at https://github.com/Swt623/Probiotic_competition.git.

## References

1. Hayden, M. K. et al. Reduction in acquisition of vancomycin-resistant enterococcus after enforcement of routine environmental cleaning measures. Clin. Infect. Dis. an Off. Publ. Infect. Dis. Soc. Am. 42, 1552–1560 (2006).

2. Sitzlar, B. et al. An environmental disinfection odyssey: evaluation of sequential interventions to improve disinfection of Clostridium difficile isolation rooms. Infect. Control Hosp. Epidemiol. 34, 459–465 (2013).

3. Quinn, M. M. et al. Cleaning and disinfecting environmental surfaces in health care: Toward an integrated framework for infection and occupational illness prevention. Am. J. Infect. Control 43, 424–434 (2015).

4. Velazquez, S. et al. From one species to another: A review on the interaction between chemistry and microbiology in relation to cleaning in the built environment. Indoor Air 29, 880–894 (2019).

5. Barber, O. W. & Hartmann, E. M. Benzalkonium chloride: A systematic review of its environmental entry through wastewater treatment, potential impact, and mitigation strategies. Crit. Rev. Environ. Sci. Technol. 0, 1–30 (2021).

6. Hora, P. I., Pati, S. G., McNamara, P. J. & Arnold, W. A. Increased Use of Quaternary Ammonium Compounds during the SARS-CoV-2 Pandemic and Beyond: Consideration of Environmental Implications. Environ. Sci. Technol. Lett. 7, 622–631 (2020).

7. Vandini, A. et al. Hard Surface Biocontrol in Hospitals Using Microbial-Based Cleaning Products. PLoS One 9, 1–13 (2014).

8. Spök, A., Arvanitakis, G. & McClung, G. Status of microbial based cleaning products in statutory regulations and ecolabelling in Europe, the USA, and Canada. Food Chem. Toxicol. 116, 10–19 (2018).

9. Caselli, E. et al. Impact of a probiotic-based cleaning intervention on the microbiota ecosystem of the hospital surfaces: Focus on the resistome remodulation. PLoS ONE vol. 11 (2016).

10. D’Accolti, M. et al. Potential of an eco-sustainable probiotic-cleaning formulation in reducing infectivity of enveloped viruses. Viruses vol. 13 (2021).

11. Stone, W., Tolmay, J., Tucker, K. & Wolfaardt, G. M. Disinfectant, soap or probiotic cleaning? Surface microbiome diversity and biofilm competitive exclusion. Microorganisms 8, 1–19 (2020).

12. Kelley, S. T. & Gilbert, J. A. Studying the microbiology of the indoor environment. Genome Biol. 14, 1–9 (2013).

13. Hu, J. et al. Impacts of indoor surface finishes on bacterial viability. Indoor Air 29, 551– 562 (2019).

14. Kokubo, M. et al. Relationship between the microbiome and indoor temperature/humidity in a traditional japanese house with a thatched roof in kyoto, japan. Diversity vol. 13 (2021).

15. Da Fonseca, T. A. P., Pessôa, R., Felix, A. C. & Sanabani, S. S. Diversity of bacterial communities on four frequently used surfaces in a large Brazilian teaching hospital. Int. J. Environ. Res. Public Health 13, 1–11 (2016).

16. Christoff, A. P. et al. One year cross-sectional study in adult and neonatal intensive care units reveals the bacterial and antimicrobial resistance genes profiles in patients and hospital surfaces. PLoS ONE vol. 15 (2020).

17. Dziri, R. et al. Characterization of extended-spectrum β-lactamase (ESBL)-producing Klebsiella, Enterobacter, and Citrobacter obtained in environmental samples of a Tunisian hospital. Diagn. Microbiol. Infect. Dis. 86, 190–193 (2016).

18. Pochtovyi, A. A. et al. Contamination of hospital surfaces with bacterial pathogens under the current COVID-19 outbreak. International Journal of Environmental Research and Public Health vol. 18 (2021).

19. Wei, L. et al. Spread of Carbapenem-Resistant Klebsiella pneumoniae in an Intensive Care Unit: A Whole-Genome Sequence-Based Prospective Observational Study. Microbiology Spectrum vol. 9 (2021).

20. Labi, A. K. et al. High carriage rates of multidrug-resistant gram-negative bacteria in neonatal intensive care units from Ghana. Open Forum Infect. Dis. 7, (2020).

21. Sereia, A. F. R. et al. Healthcare-Associated Infections-Related Bacteriome and Antimicrobial Resistance Profiling: Assessing Contamination Hotspots in a Developing Country Public Hospital. Front. Microbiol. 12, 1–18 (2021).

22. Al-Hamad, A. et al. Molecular characterization of clinical and environmental carbapenem resistant Acinetobacter baumannii isolates in a hospital of the Eastern Region of Saudi Arabia. J. Infect. Public Health 13, 632–636 (2020).

23. Khalilzadegan, S. et al. Beta-Lactamase Encoded Genes blaTEM and blaCTX Among Acinetobacter baumannii Species Isolated From Medical Devices of Intensive Care Units in Tehran Hospitals. Jundishapur J. Microbiol. 9, e14990 (2016).

24. Rose, M., Landman, D. & Quale, J. Are community environmental surfaces near hospitals reservoirs for gram-negative nosocomial pathogens? Am. J. Infect. Control 42, 346–348 (2014).

25. Elkady, M. A., Bakr, W. M. K., Ghazal, H. & Omran, E. A. Role of environmental surfaces and hands of healthcare workers in perpetuating multi-drug-resistant pathogens in a neonatal intensive care unit. European Journal of Pediatrics vol. 181 619–628 (2022).

26. Bouguenoun, W. et al. Molecular epidemiology of environmental and clinical carbapenemase-producing Gram-negative bacilli from hospitals in Guelma, Algeria: Multiple genetic lineages and first report of OXA-48 in Enterobacter cloacae. J. Glob. Antimicrob. Resist. 7, 135–140 (2016).

27. Brooks, B. et al. Strain-resolved analysis of hospital rooms and infants reveals overlap between the human and room microbiome. Nat. Commun. 8, 1–7 (2017).

28. Brooks, B. et al. Microbes in the neonatal intensive care unit resemble those found in the gut of premature infants. Microbiome 2, 1–16 (2014).

29. Cason, C. et al. Microbial contamination in hospital environment has the potential to colonize preterm newborns’ nasal cavities. Pathogens vol. 10 (2021).

30. Ng, D. H. L. et al. Environmental colonization and onward clonal transmission of carbapenem-resistant Acinetobacter baumannii (CRAB) in a medical intensive care unit: The case for environmental hygiene. Antimicrobial Resistance and Infection Control vol. 7 (2018).

31. Chen, L. F. et al. A prospective study of transmission of Multidrug-Resistant Organisms (MDROs) between environmental sites and hospitalized patients - The TransFER study. Infect. Control Hosp. Epidemiol. 40, 47–52 (2019).

32. Nguyen, T. N. T. et al. Emerging carbapenem-resistant Klebsiella pneumoniae sequence type 16 causing multiple outbreaks in a tertiary hospital in southern Vietnam. Microb. Genomics 7, (2021).

33. Senok, A. et al. Genetic relatedness of clinical and environmental acinetobacter baumanii isolates from an intensive care unit outbreak. J. Infect. Dev. Ctries. 9, 665–669 (2015).

34. Shen, J., McFarland, A. G., Young, V. B., Hayden, M. K. & Hartmann, E. M. Toward Accurate and Robust Environmental Surveillance Using Metagenomics. Front. Genet. 12, 1–8 (2021).

35. Song, W. et al. Modified CHROMagar Acinetobacter medium for direct detection of multidrug-resistant Acinetobacter strains in nasal and rectal swab samples. Ann. Lab. Med. 33, 193–195 (2013).

36. Bankevich, A. et al. SPAdes: a new genome assembly algorithm and its applications to single-cell sequencing. J. Comput. Biol. 19, 455–477 (2012).

37. Seemann, T. Prokka: rapid prokaryotic genome annotation. Bioinformatics 30, 2068–2069 (2014).

38. Jia, B. et al. CARD 2017: expansion and model-centric curation of the comprehensive antibiotic resistance database. Nucleic Acids Res. 45, D566–D573 (2017).

39. Uritskiy, G. V, DiRuggiero, J. & Taylor, J. MetaWRAP—a flexible pipeline for genome-resolved metagenomic data analysis. Microbiome 6, 158 (2018).

40. Parks, D. H., Imelfort, M., Skennerton, C. T., Hugenholtz, P. & Tyson, G. W. CheckM: assessing the quality of microbial genomes recovered from isolates, single cells, and metagenomes. Genome Res. 25, 1043–1055 (2015).

41. Chaumeil, P.-A., Mussig, A. J., Hugenholtz, P. & Parks, D. H. GTDB-Tk: a toolkit to classify genomes with the Genome Taxonomy Database. Bioinformatics 36, 1925–1927 (2020).

42. Kopylova, E., Noé, L. & Touzet, H. SortMeRNA: fast and accurate filtering of ribosomal RNAs in metatranscriptomic data. Bioinformatics 28, 3211–3217 (2012).

43. Kim, D., Langmead, B. & Salzberg, S. L. HISAT: a fast spliced aligner with low memory requirements. Nat. Methods 12, 357–360 (2015).

44. Liao, Y., Smyth, G. K. & Shi, W. featureCounts: an efficient general purpose program for assigning sequence reads to genomic features. Bioinformatics 30, 923–930 (2014).

45. Love, M. I., Huber, W. & Anders, S. Moderated estimation of fold change and dispersion for RNA-seq data with DESeq2. Genome Biol. 15, 550 (2014).

46. Kanehisa, M., Sato, Y. & Morishima, K. BlastKOALA and GhostKOALA: KEGG Tools for Functional Characterization of Genome and Metagenome Sequences. J. Mol. Biol. 428, 726–731 (2016).

47. Yu, G., Wang, L.-G., Han, Y. & He, Q.-Y. clusterProfiler: an R package for comparing biological themes among gene clusters. OMICS 16, 284–287 (2012).

48. Lin, K., Vikesland, P. J. & Isaacman-Vanwertz, G. Viability of Viruses in Suspended Aerosols and Stationary Droplets as a Function of Relative Humidity and Media Composition. (2020).

49. Farrow, J. M., Wells, G., Palethorpe, S., Adams, M. D. & Pesci, E. C. CsrA supports both environmental persistence and host-associated growth of acinetobacter baumannii. Infection and Immunity vol. 88 (2020).

50. Krasauskas, R., Skerniškytė, J., Armalytė, J. & Sužiedėlienė, E. The role of Acinetobacter baumannii response regulator BfmR in pellicle formation and competitiveness via contact-dependent inhibition system. BMC Microbiol. 19, 1–12 (2019).

51. Fenollar, F. & Mediannikov, O. Emerging infectious diseases in Africa in the 21st century. New microbes new Infect. 26, S10–S18 (2018).

52. Lin, K. & Marr, L. C. Humidity-Dependent Decay of Viruses, but Not Bacteria, in Aerosols and Droplets Follows Disinfection Kinetics. Environ. Sci. Technol. 54, 1024– 1032 (2020).

53. Antunes, L. C. S., Visca, P. & Towner, K. J. Acinetobacter baumannii: Evolution of a global pathogen. Pathog. Dis. 71, 292–301 (2014).

54. Avrani, S., Bolotin, E., Katz, S. & Hershberg, R. Rapid Genetic Adaptation during the First Four Months of Survival under Resource Exhaustion. Mol. Biol. Evol. 34, 1758– 1769 (2017).

55. Behringer, M. G. et al. Escherichia coli cultures maintain stable subpopulation structure during long-term evolution. Proc. Natl. Acad. Sci. U. S. A. 115, E4642–E4650 (2018).

56. Martin, R. M. & Bachman, M. A. Colonization, infection, and the accessory genome of Klebsiella pneumoniae. Front. Cell. Infect. Microbiol. 8, 1–15 (2018).

57. Stubbendieck, R. M. & Straight, P. D. Multifaceted Interfaces of Bacterial Competition. J. Bacteriol. 198, 2145–2155 (2016).

58. Butaitė, E., Baumgartner, M., Wyder, S. & Kümmerli, R. Siderophore cheating and cheating resistance shape competition for iron in soil and freshwater Pseudomonas communities. Nat. Commun. 8, 414 (2017).

59. Hsieh, P.-F., Lu, Y.-R., Lin, T.-L., Lai, L.-Y. & Wang, J.-T. Klebsiella pneumoniae Type VI Secretion System Contributes to Bacterial Competition, Cell Invasion, Type-1 Fimbriae Expression, and In Vivo Colonization. J. Infect. Dis. 219, 637–647 (2019).

60. Struve, C., Bojer, M. & Krogfelt, K. A. Characterization of Klebsiella pneumoniae type 1 fimbriae by detection of phase variation during colonization and infection and impact on virulence. Infect. Immun. 76, 4055–4065 (2008).

61. Itoh, Y. et al. Roles of pgaABCD genes in synthesis, modification, and export of the Escherichia coli biofilm adhesin poly-beta-1,6-N-acetyl-D-glucosamine. J. Bacteriol. 190, 3670–3680 (2008).

62. Wang, X., Preston, J. F. 3rd & Romeo, T. The pgaABCD locus of Escherichia coli promotes the synthesis of a polysaccharide adhesin required for biofilm formation. J. Bacteriol. 186, 2724–2734 (2004).

63. Sharma, A., Sharma, R., Bhattacharyya, T., Bhando, T. & Pathania, R. Fosfomycin resistance in Acinetobacter baumannii is mediated by efflux through a major facilitator superfamily (MFS) transporter-AbaF. J. Antimicrob. Chemother. 72, 68–74 (2017).

64. Beale, J., Lee, S. Y., Iwata, S. & Beis, K. Structure of the aliphatic sulfonate-binding protein SsuA from Escherichia coli. Acta Crystallogr. Sect. F Struct. Biol. Cryst. Commun. 66, 391–396 (2010).

65. van der Ploeg, J. R. et al. Identification of sulfate starvation-regulated genes in Escherichia coli: a gene cluster involved in the utilization of taurine as a sulfur source. J. Bacteriol. 178, 5438–5446 (1996).

66. Khleifat, K. M. Biodegradation of sodium lauryl ether sulfate (SLES) by two different bacterial consortia. Curr. Microbiol. 53, 444–448 (2006).

67. van der Ploeg, J. R., Eichhorn, E. & Leisinger, T. Sulfonate-sulfur metabolism and its regulation in Escherichia coli. Arch. Microbiol. 176, 1–8 (2001).

68. Dhouib, A., Hamad, N., Hassaıri, I. & Sayadi, S. Degradation of anionic surfactants by Citrobacter braakii. Process Biochem. 38, 1245–1250 (2003).

69. Hu, J. & Hartmann, E. M. Anthropogenic chemicals and their impacts on microbes living in buildings. Microb. Biotechnol. 14, 798–802 (2021).

70. Subramanian, A. et al. Gene set enrichment analysis: a knowledge-based approach for interpreting genome-wide expression profiles. Proc. Natl. Acad. Sci. U. S. A. 102, 15545– 15550 (2005).

71. Mootha, V. K. et al. PGC-1alpha-responsive genes involved in oxidative phosphorylation are coordinately downregulated in human diabetes. Nat. Genet. 34, 267–273 (2003).

72. De Araujo, C., Balestrino, D., Roth, L., Charbonnel, N. & Forestier, C. Quorum sensing affects biofilm formation through lipopolysaccharide synthesis in Klebsiella pneumoniae. Res. Microbiol. 161, 595–603 (2010).

73. Falagas, M. E. & Makris, G. C. Probiotic bacteria and biosurfactants for nosocomial infection control: a hypothesis. J. Hosp. Infect. 71, 301–306 (2009).

74. Al-Marzooq, F. et al. Can probiotic cleaning solutions replace chemical disinfectants in dental clinics? Eur. J. Dent. 12, 532–539 (2018).

75. Caselli, E. et al. Impact of a Probiotic-Based Cleaning Intervention on the Microbiota Ecosystem of the Hospital Surfaces: Focus on the Resistome Remodulation. PLoS One 11, 1–19 (2016).

76. Caselli, E. et al. Impact of a probiotic-based hospital sanitation on antimicrobial resistance and HAI-associated antimicrobial consumption and costs: a multicenter study. Infect. Drug Resist. 12, 501–510 (2019).

77. Suez, J., Zmora, N., Segal, E. & Elinav, E. The pros, cons, and many unknowns of probiotics. Nat. Med. 25, 716–729 (2019).

78. Andualem, Z., Gizaw, Z., Bogale, L. & Dagne, H. Indoor bacterial load and its correlation to physical indoor air quality parameters in public primary schools. Multidiscip. Respir. Med. 14, 2 (2019).

79. Elshaghabee, F. M. F., Rokana, N., Gulhane, R. D., Sharma, C. & Panwar, H. Bacillus As Potential Probiotics: Status, Concerns, and Future Perspectives. Front. Microbiol. 8, 1490 (2017).

80. Elisashvili, V., Kachlishvili, E. & Chikindas, M. L. Recent Advances in the Physiology of Spore Formation for Bacillus Probiotic Production. Probiotics Antimicrob. Proteins 11, 731–747 (2019).

