## Supplemental Info for "Clinically Relevant Pathogens on Surfaces Display Differences in Survival and Transcriptomic Response in Relation to Probiotic and Traditional Cleaning Strategies"

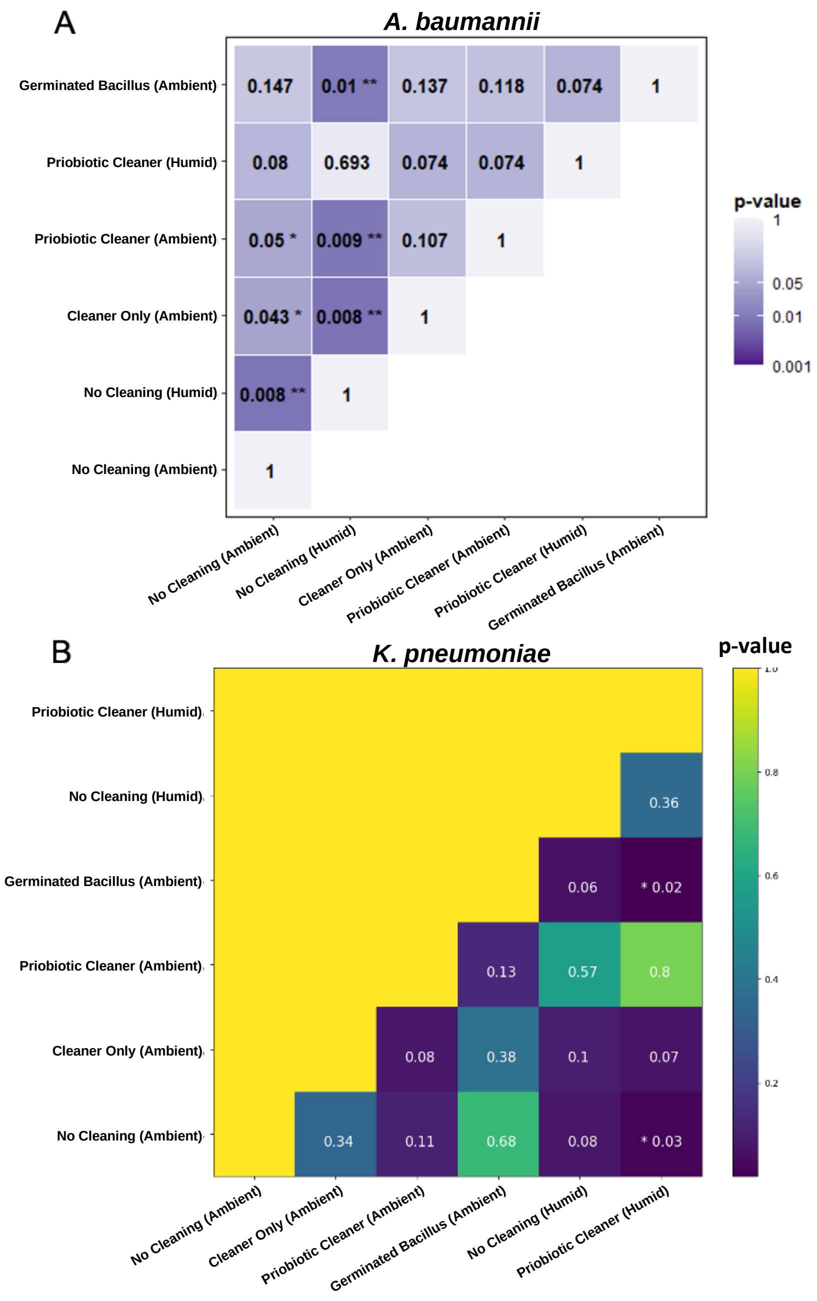


**Figure S1.** p-values of *A. baumannii* (A) and *K. pneumoniae* (B) viable CFU counts among samples collected under two temperature/humidity conditions and four cleaning scenarios. Statistical testing was conducted using Welch's paired t-test. * p<0.05, ** p< 0.01, and ***p < 0.001.

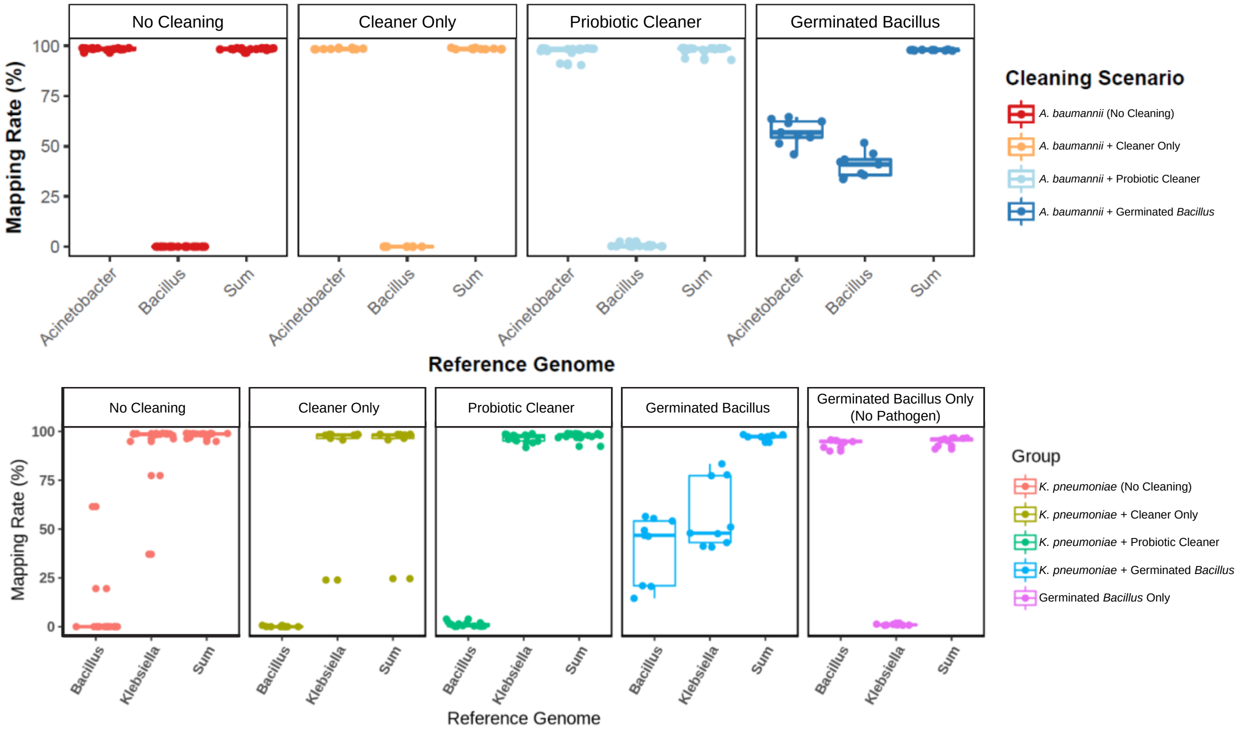


**Figure S2.** Percentages of reads mapped onto *Bacillus* spp. and either *A. baumannii* (top) or *K. pneumoniae* (bottom) genomes for samples collected from four surface cleaning scenarios.

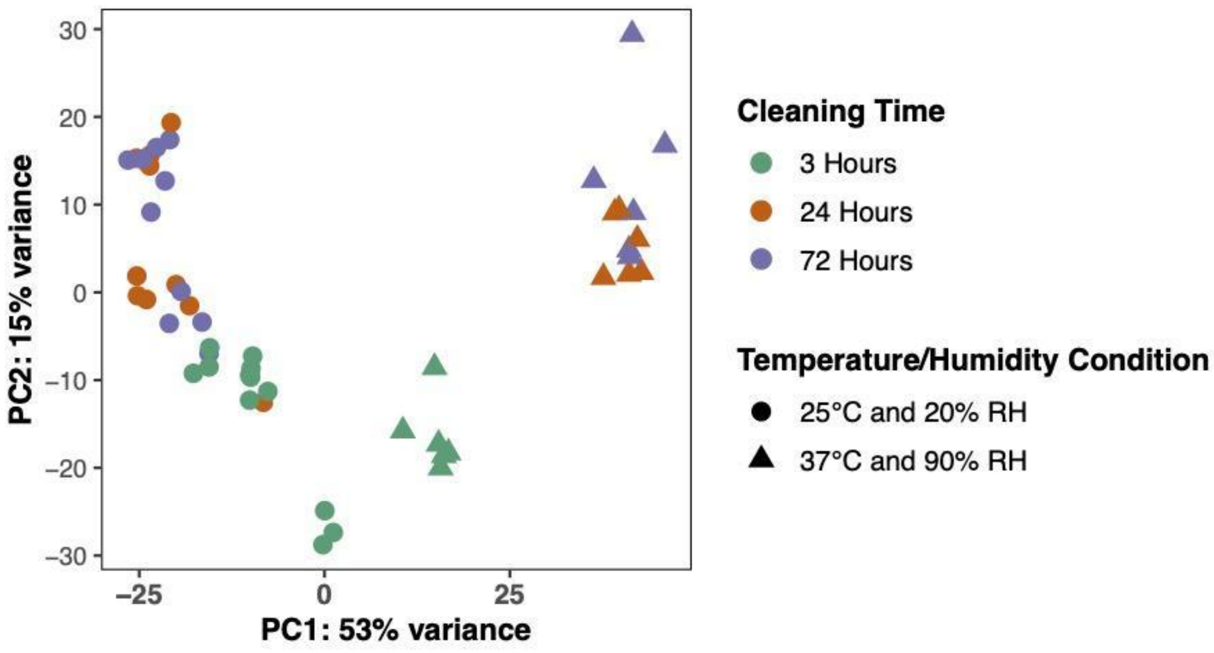


**Figure S3.** Principal components analysis on variance stabilized *K. pneumoniae* read counts. Samples were color-coded based on cleaning time.

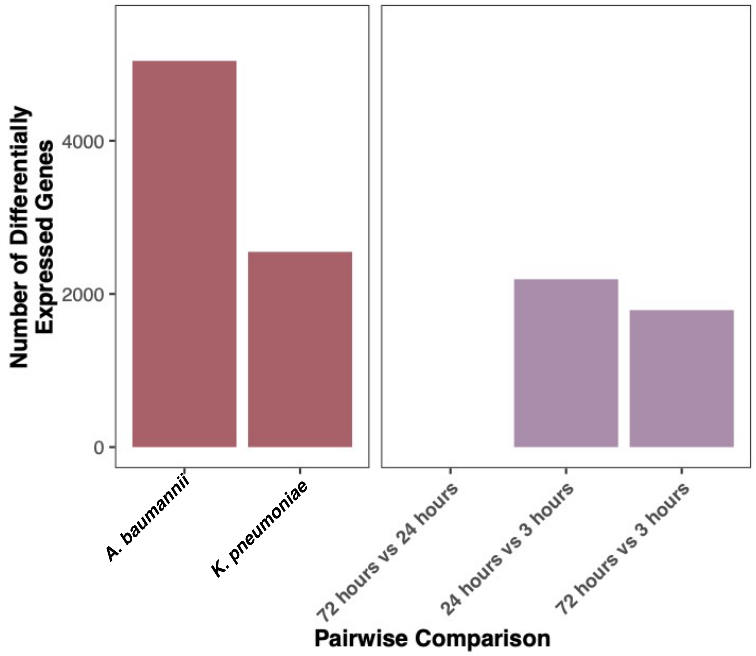


**Figure S4**. Number of differentially expressed genes for germinated *Bacillus* reads. The *Bacillus* transcriptome was compared in the presence of *A. baumannii* or *K. pneumoniae* using vegetative *Bacillus* without pathogen as the reference level. Comparison was also conducted between different time points.

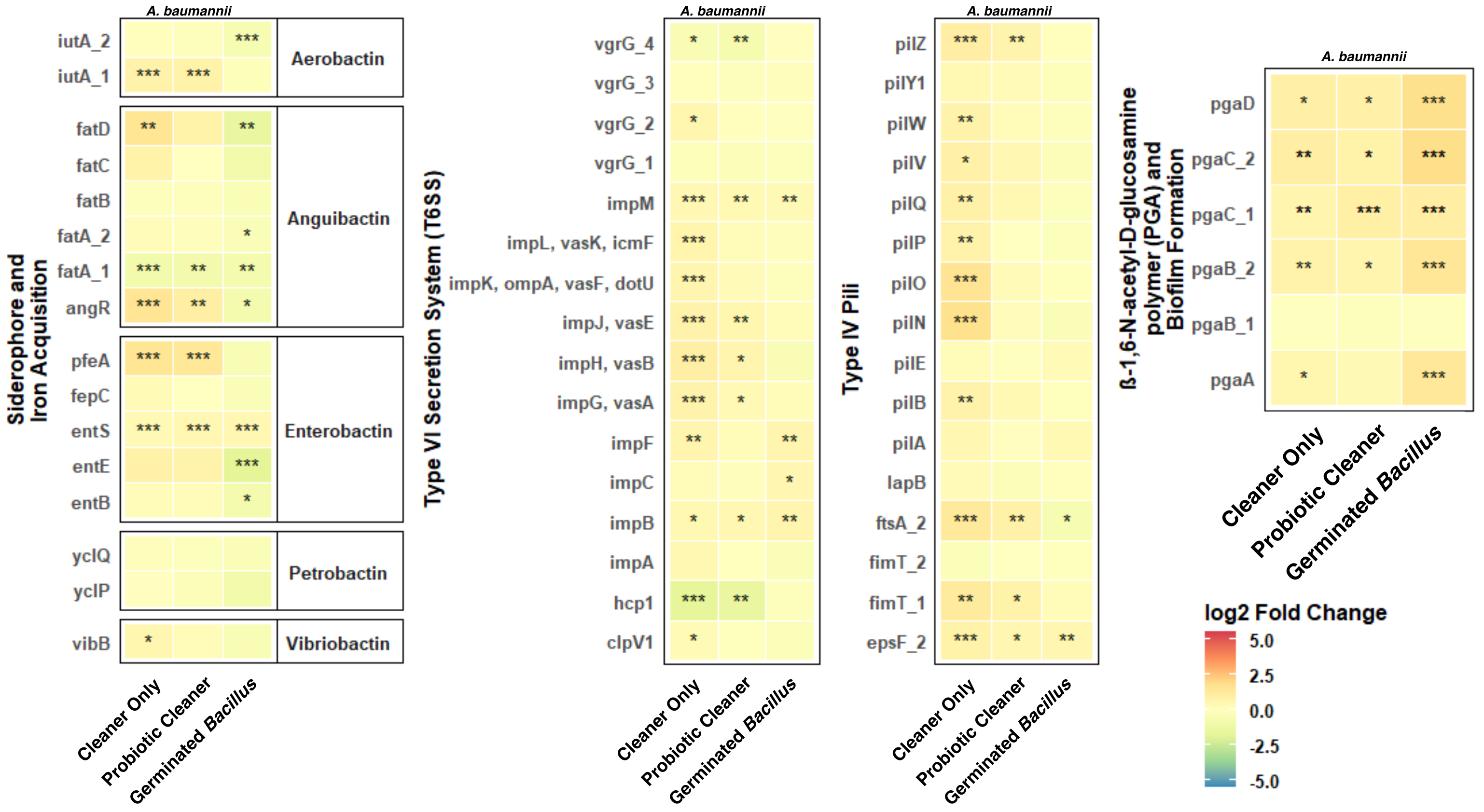


**Figure S5.** Log2 fold change of genes associated with iron acquisition, Type VI Secretion System (T6SS), Type VI Pili, and biofilm poly-beta-1,6-N-acetyl-D-glucosamine (PGA) production in samples cleaned with Cleaner Only, Probiotic Cleaner, and Germinated *Bacillus* compared to No Cleaner samples (*A. baumannii* only). Statistically significant log2 fold changes were determined by DESeq2 with Benjamini-Hochberg corrected p-value ≤ 0.05. * p<0.05, ** p< 0.01, and ***p < 0.001.

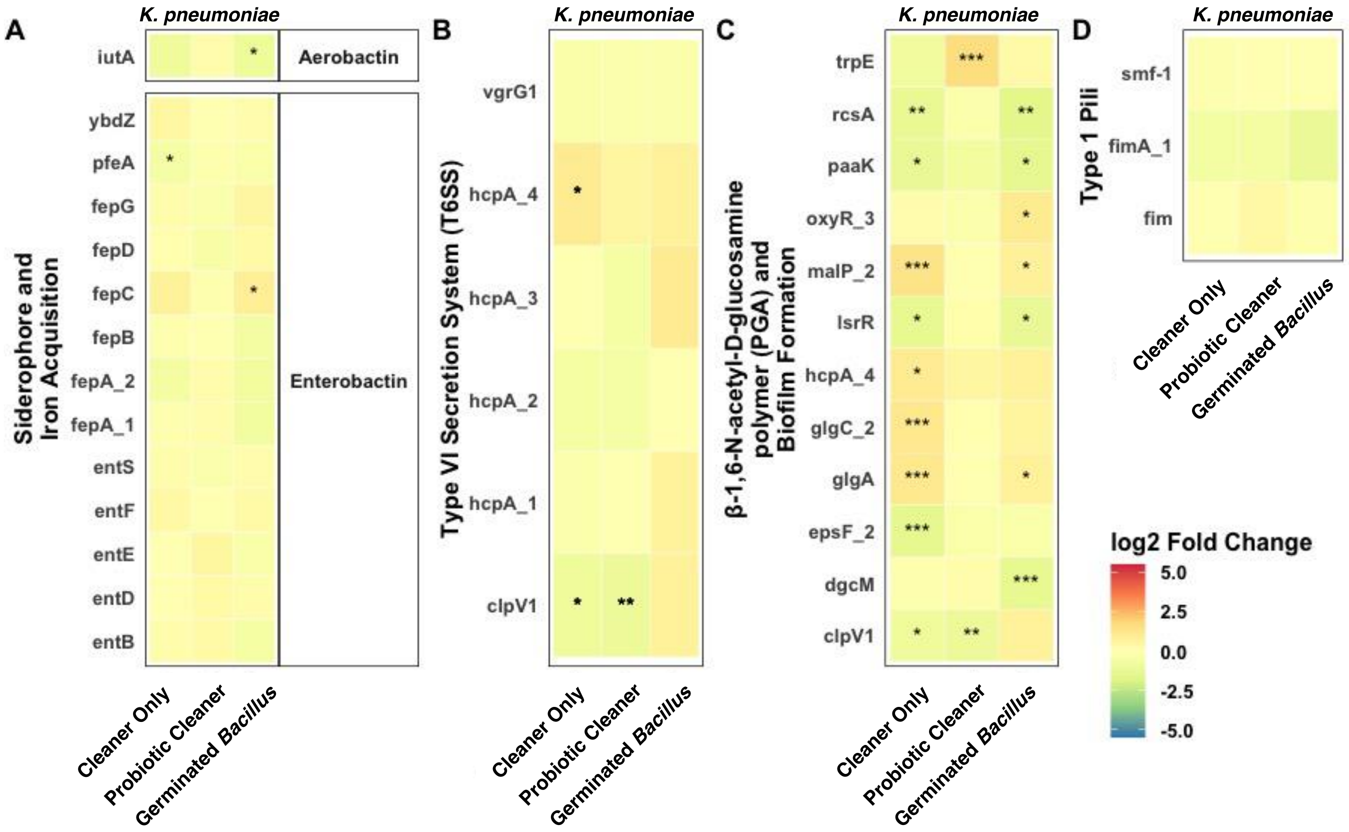


**Figure S6.** Log2 fold change of genes associated with iron acquisition, Type VI Secretion System (T6SS), biofilm poly-beta-1,6-N-acetyl-D-glucosamine (PGA) production, and Type 1 Pili in samples cleaned with Cleaner Only, Probiotic Cleaner, and Germinated *Bacillus* compared to No Cleaner samples (*K. pneumoniae* only). Statistically significant log2 fold changes were determined by DESeq2 with Benjamini-Hochberg corrected p-value ≤ 0.05. Some genes have Benjamini-Hochberg corrected p-value ≤ 0.05 but log2 fold change < 1. * p<0.05, ** p< 0.01, and ***p < 0.001.

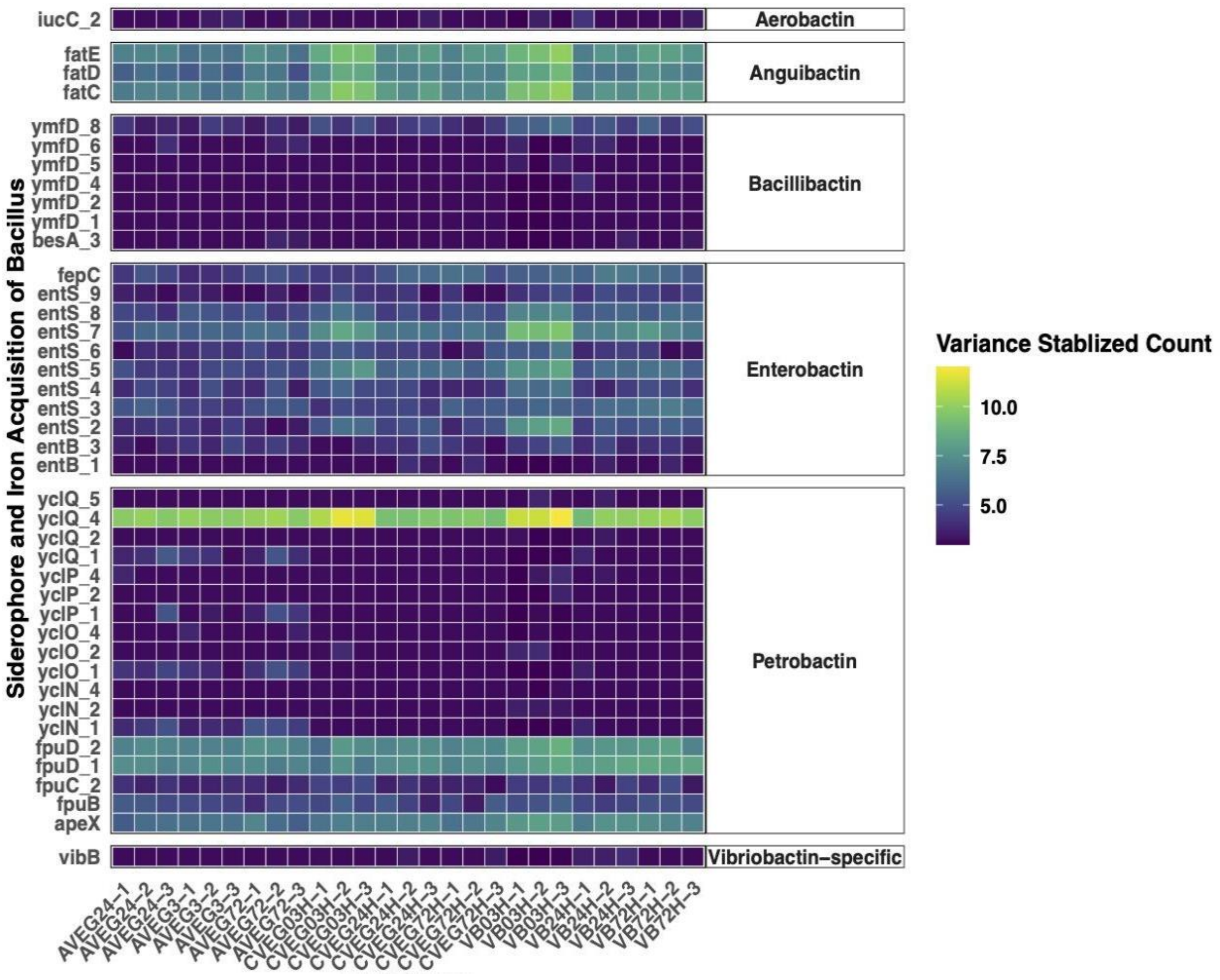


**Figure S7.** Expression of genes associated with siderophore and iron acquisition in vegetative *Bacillus* spp. samples in the presence of *A. baumannii* (AVEG), *K. pneumoniae* (CVEG), and in the absence of pathogen (VB). Read counts belonging to Bacillus *spp*. were transformed using variance stabilization to enable between-sample comparisons.
